# *In-silico* evaluation of Mirror repeats in some selected genes of *Candida albicans*

**DOI:** 10.1101/2024.01.05.574287

**Authors:** Barkha Sehrawat, Priya Yadav, Mustak Sarjeet, Sakshi Yadav, Vidhi Yadav, Nupur Goyal, Parvej Alam, Meghali Ahlawat, Sandeep Yadav

## Abstract

All cellular processes in a living cell are controlled by its genetic material. DNA in majority of the domains acts as a regulatory molecule by controlling various vital functions. The genetic makeup of DNA or RNA (in viruses) is unique in all the domains. Their uniqueness is determined by the presence of various types of repetitive patterns of bases. These includes inverted repeats, tandem repeats, VNTR’s, palindromes etc. Among many repetitive pattern types, Mirror repeats (MR) found to be dispersed throughout the genes or genomes. These sequences are associated with various functional features like their involvement in H-DNA formation, in replication & transcription, nervous system related diseases development etc. The major focus of this investigation is to identify MR sequences from some selected genes of *Candida albicans* using a bioinformatics based pipeline. The approach refers to as FPCB which utilized some manual steps to extract out MR sequence from any targeted gene or genome. The current study find out that the identified Mirror repeats found to be dispersed throughout the selected genes along with variable length. Among them the maximum & minimum MR sequences were reported in the gene FAS2 (108) & HIS1 (15) respectively. The present study will be helpful to provide a new insight for molecular as well as computational based studies of *Candida albicans* as well as other related fungal groups.

## Introduction

What makes a specific organism different from one another is their genetic material which is acts as a blueprint of life (1). Genetic material could be either DNA (in case of bacteria, animals, fungi, plant etc.) or RNA (in case of some viruses like HIV, flu virus like corona virus etc.) It is composed of repeated nucleotides units which also form a unique pattern of sequences that makes all organisms different from others. The sequence pattern which is composed of base pairs shows diverse group of repeated sequence (2). These repetitive sequences are differentiated into two types on the basis of their arrangement in genes or genomes. One includes tandem repeat which is present adjacent to each to other whereas another type is interspersed repeat which is scattered throughout the genome (3-4). Tandem repeat further classified into microsatellite & minisatellite DNA (5). Similarly Interspersed repeat consist of SINE & LINE sequences and retrotransposons (6-8). These repeats perform variety of functions like help in gene expression, function as evolutionary markers, makeup the promoters & enhancers parts on DNA and also associated with many diseases in human (9-11). Repeated sequences are always attracting researchers because of their unique roles at molecular level. Among these repeats a special sequence pattern refers to as Mirror repeat also reported. Mirror repeat is basically that segment of genome or gene which shares a center of symmetry, forming exact mirror image of each other on the same strand. For example, in this sequence TGACGGCATTACGGCAGT, TGACGGCAT shares center of symmetry or shows homology with rest of its part. Due to mutation in polypyrimidine and polypurine mirror repeat it causes genomic instability that results in genetic, neurological diseases (12-15). Mirror repeats have been identified in various phyla including bacteria, animal & plant viruses, as well as in human insulin gene (16-19). The present research will focus on identification of mirror repeats in some selected genes of fungi *Candida albicans* using an *in silico* technique (FPCB) (20). FPCB stands for FASTA Parallel Complement BLAST, a bioinformatics based manual pipeline to find out mirror repeats in any targeted gene or genome sequence (20-22). This study will be helpful in analysing the functioning as well as molecular prospective of mirror repeats sequence in fungal as well as other living domains.

## Materials and Methods

To identify mirror repeats in selected *Candida albicans* genes (Table-1), a bioinformatics based manual pipeline refers to as FPCB (20) were utilized. This approach can be applied on any targeted gene or genome family to extract MR sequences. It involve four different steps, in each step public domain databases were utilized which are freely accessible. The steps involve in this process (shows in Figure 1) are followings: –

**Figure 1.**
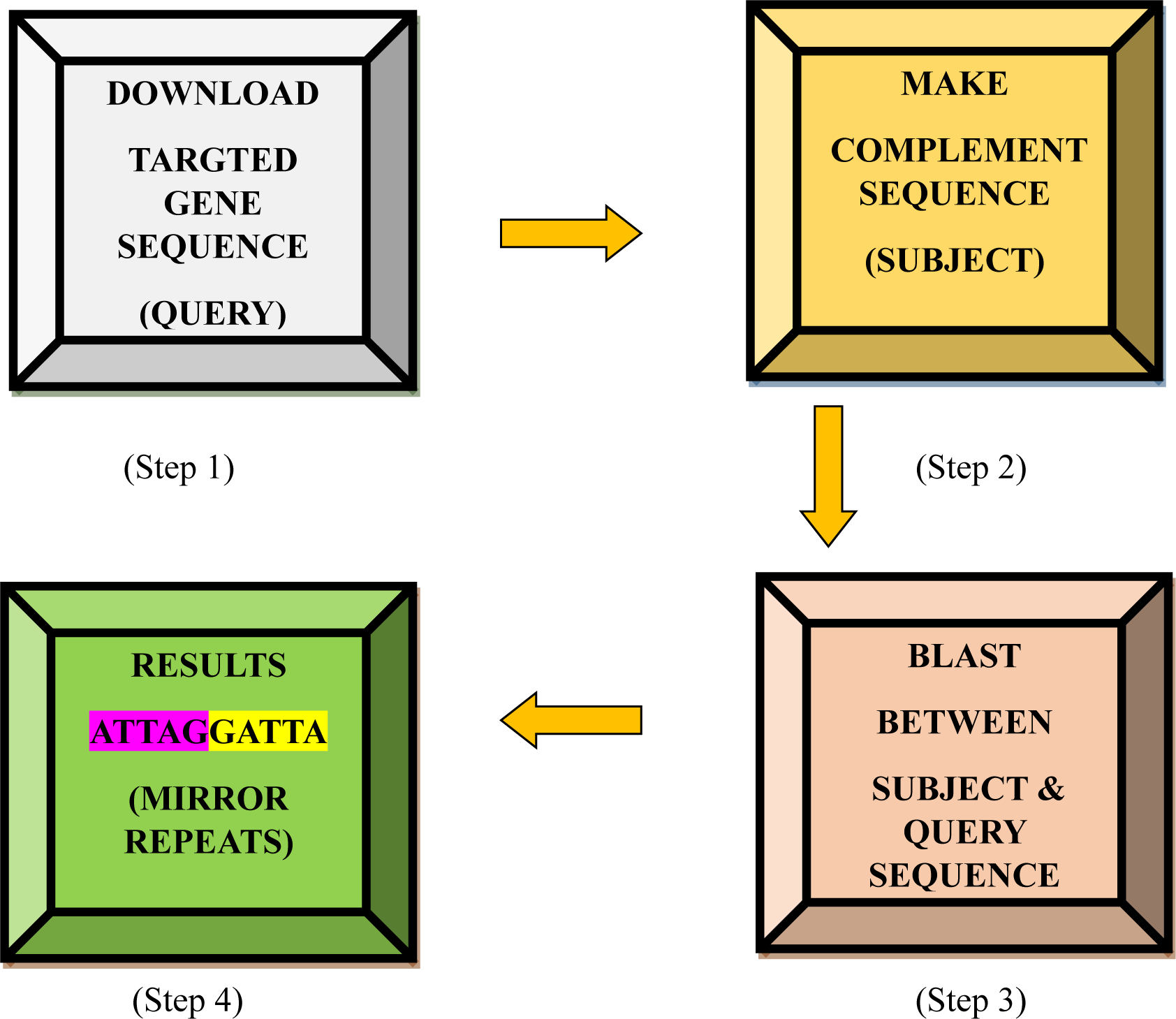
Depicted Flow Model of FPCB.

➢ The targeted gene sequence of *C. albicans* was downloaded from NCBI (23)
➢ The downloaded sequence can be processed through Reverse complement tool
➢ Subject & Query sequence align with each other using BLAST tool
➢ Result & Analysis

**Table 1.** List of Genes targeted for Mirror Repeat Identification.

| S. No. | Gene Name | Gene Functions | Gene ID |
| --- | --- | --- | --- |
| 1 | ADE 2 | Nucleotide synthesis | 3635259 |
| 2 | FAS 2 | Fatty acid synthase subunit, lipid synthesis | 3635222 |
| 3 | FTR 1 | High-affinity iron permease | 3643318 |
| 4 | HEM 3 | Haemosynthesis | 3643940 |
| 5 | HIS 1 | Amino acid biosynthesis | 3636501 |
| 6 | MIG 1 | DNA binding protein, glucose metabolism | 3637812 |
| 7 | LEU 2 | Amino acid synthesis | 3638034 |
| 8 | SNF 1 | Derepression of catabolite repression, carbon source utilization | 3642643 |
| 9 | NMT 1 | Myristoyl-CoA:protein *N*-myristoyltransferase, lipid biosynthesis | 3635624 |
| 10 | URA 3 | Nucleotide synthesis | 3636649 |

## Results

By using FPCB (a bioinformatics-based approach) we extract mirror repeats for selective genes of *Candida albicans*. The identified mirror repeats in all the 10 genes are of different lengths. The minimum length of mirror repeats were observed is of 7 nucleotide which was found in almost all genes but the maximum length of mirror repeat (MR No. 9) is of 54 nucleotide found in gene MIG 1. The graph shown below (Figure 2) represents the total number of identified mirror repeats in each selected gene. The maximum & minimum numbers of MR sequences were reported in FAS2 (108) & HIS1 (15) gene respectively.

**Figure 2.**
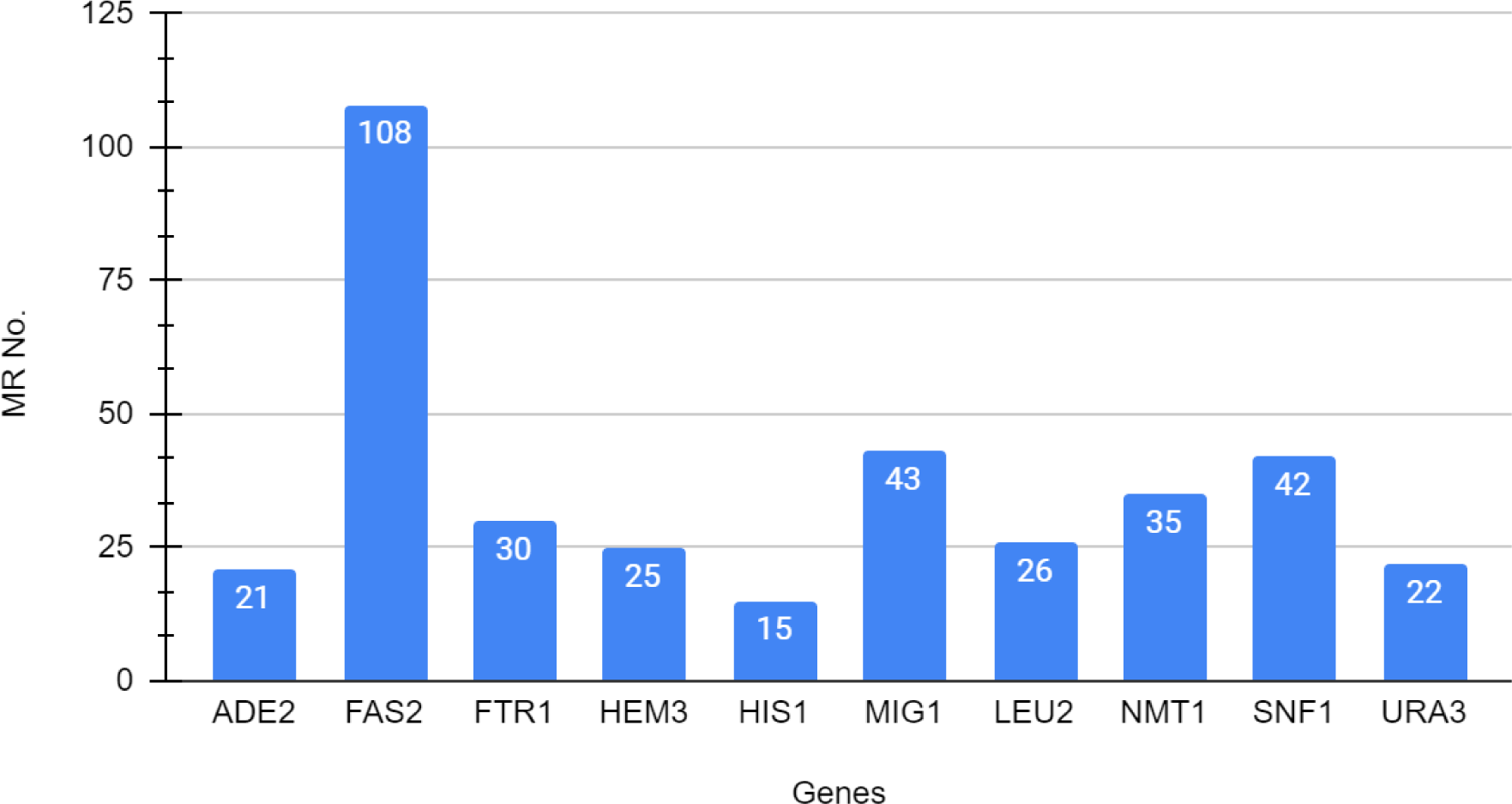
Shows graphical representation of gene wise distribution of mirror repeats in selected genes of *Candida albicans*.

The complete gene wise distribution of mirror repeats sequence in selected genes of *Candida albicans* along with their length, position in the regions and sequences are enlisted in tables below.

**Table 2.**
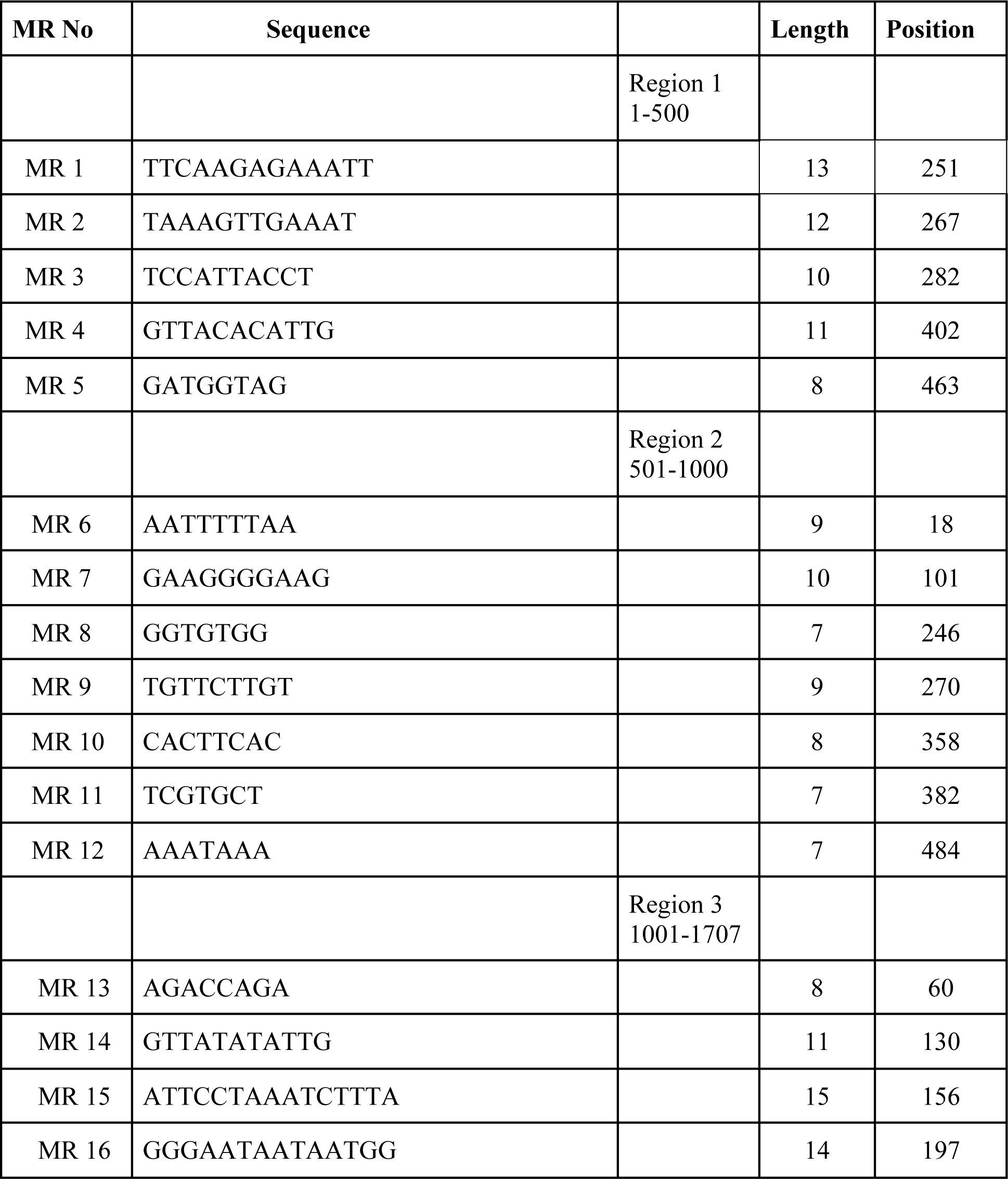
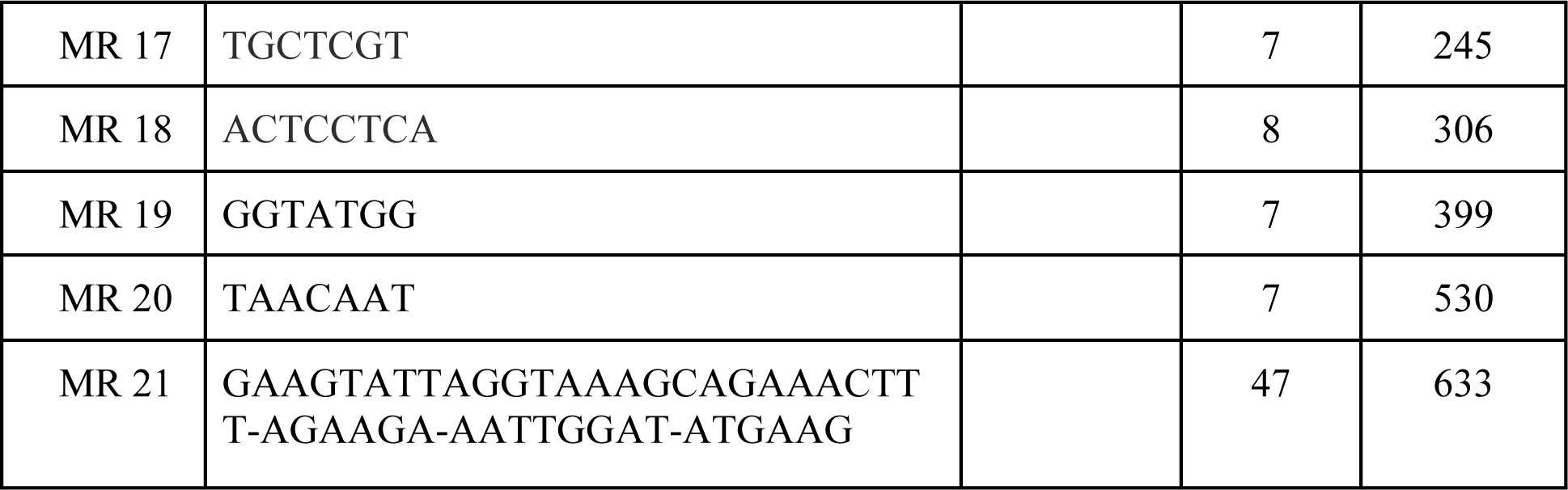
Shows Mirror repeat sequence pattern, their length & position in ADE2 gene.

**Table 3.**
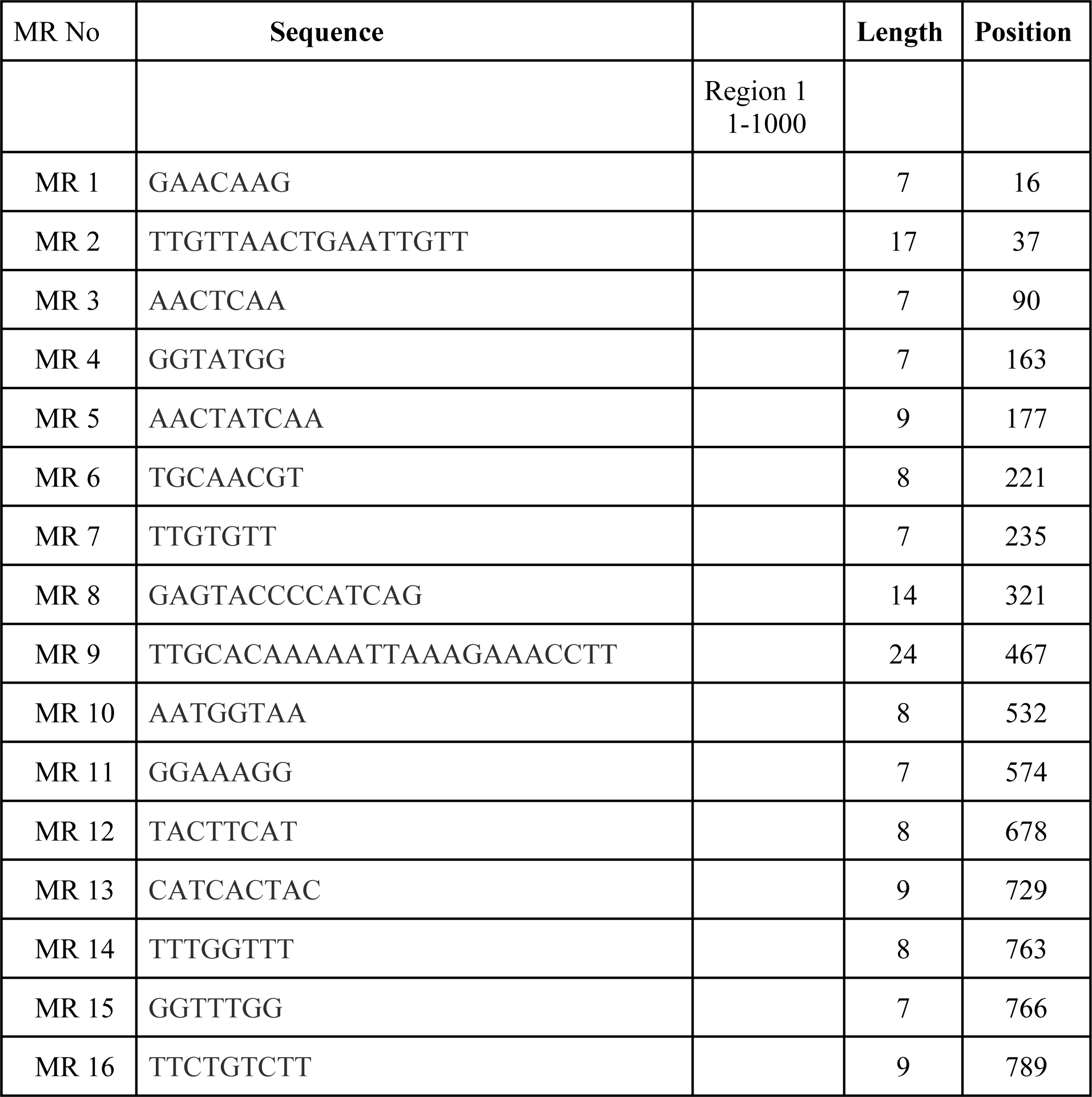
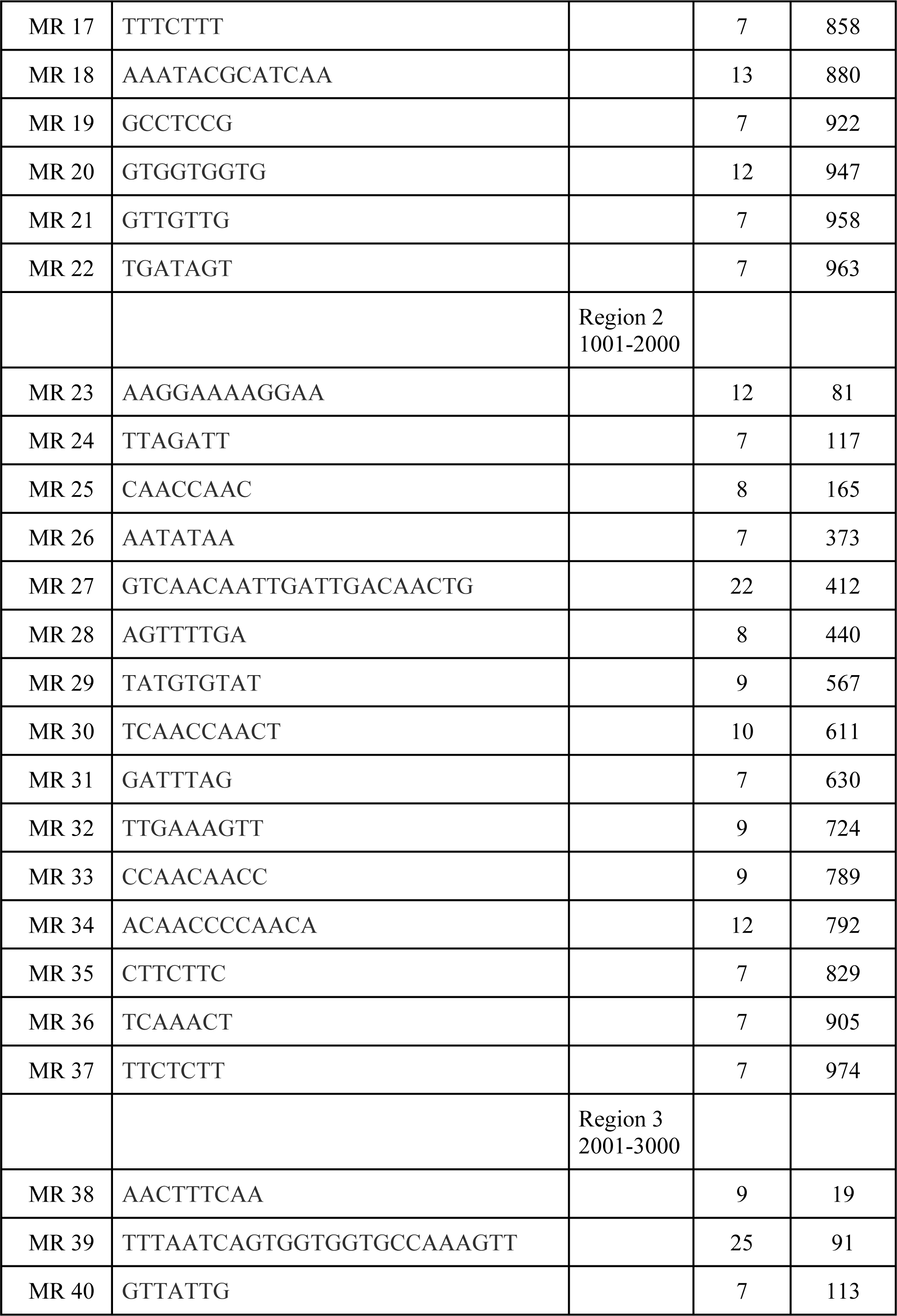
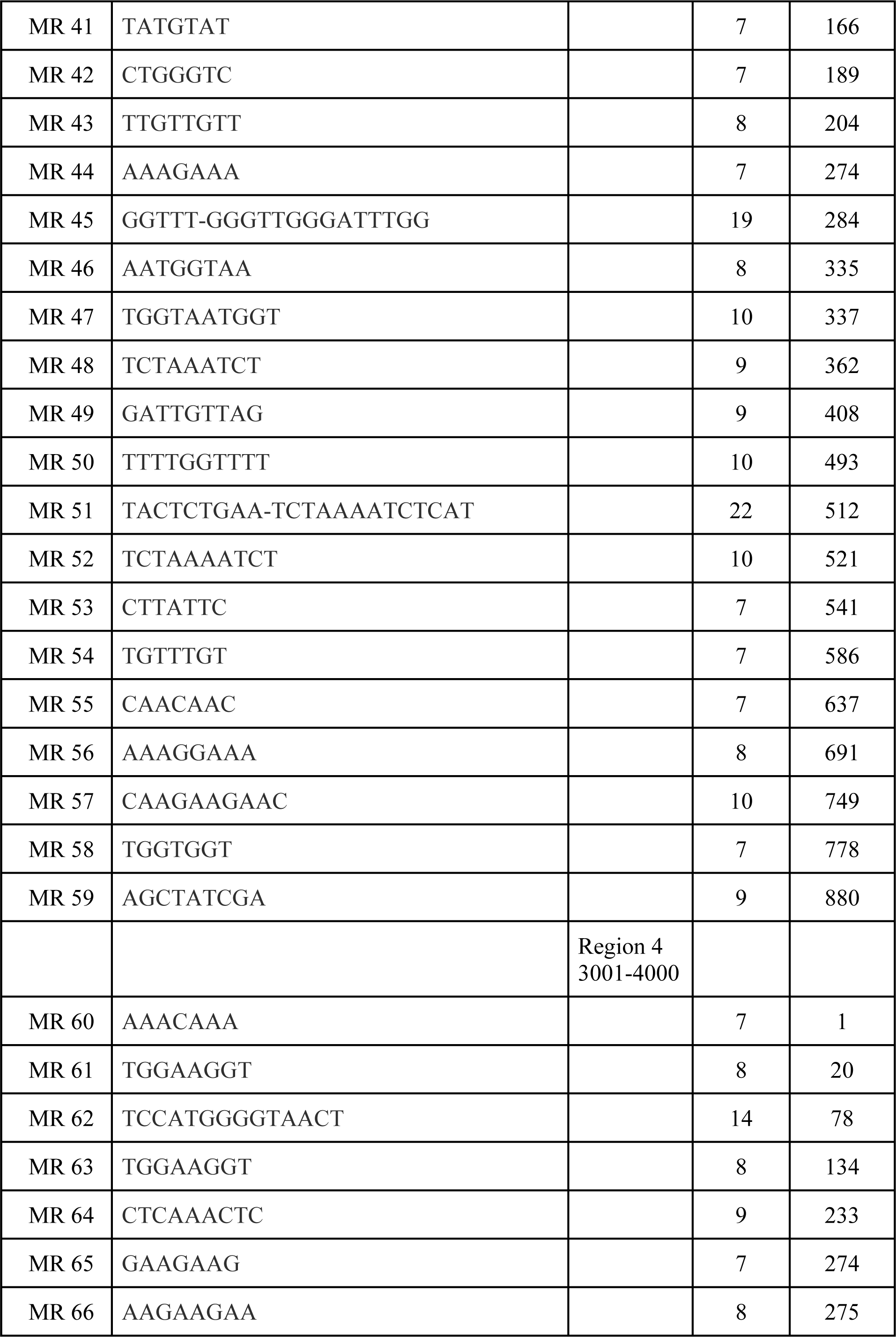
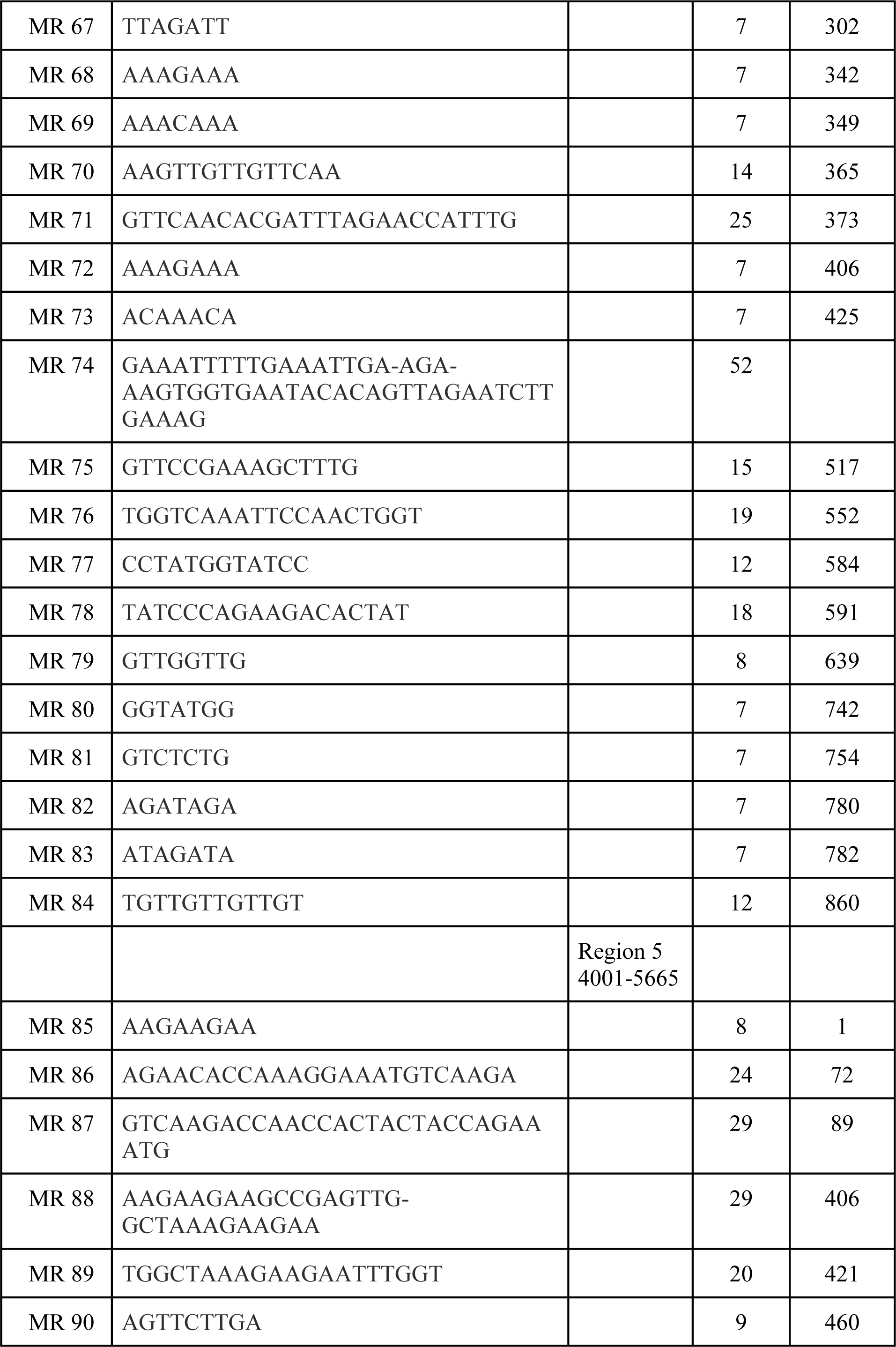
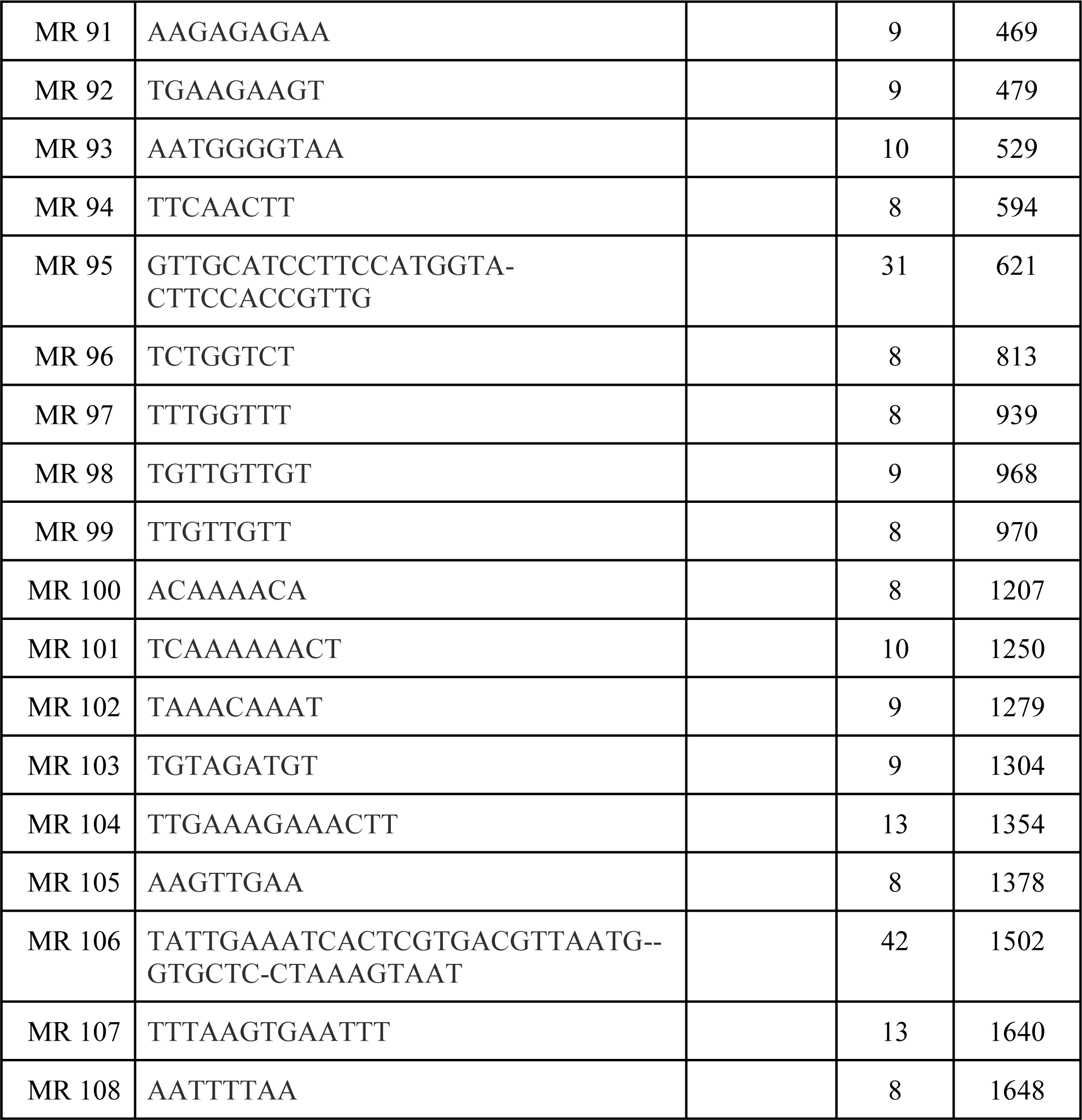
Shows Mirror repeat sequence pattern, their length & position in FAS2 gene.

**Table 4.**
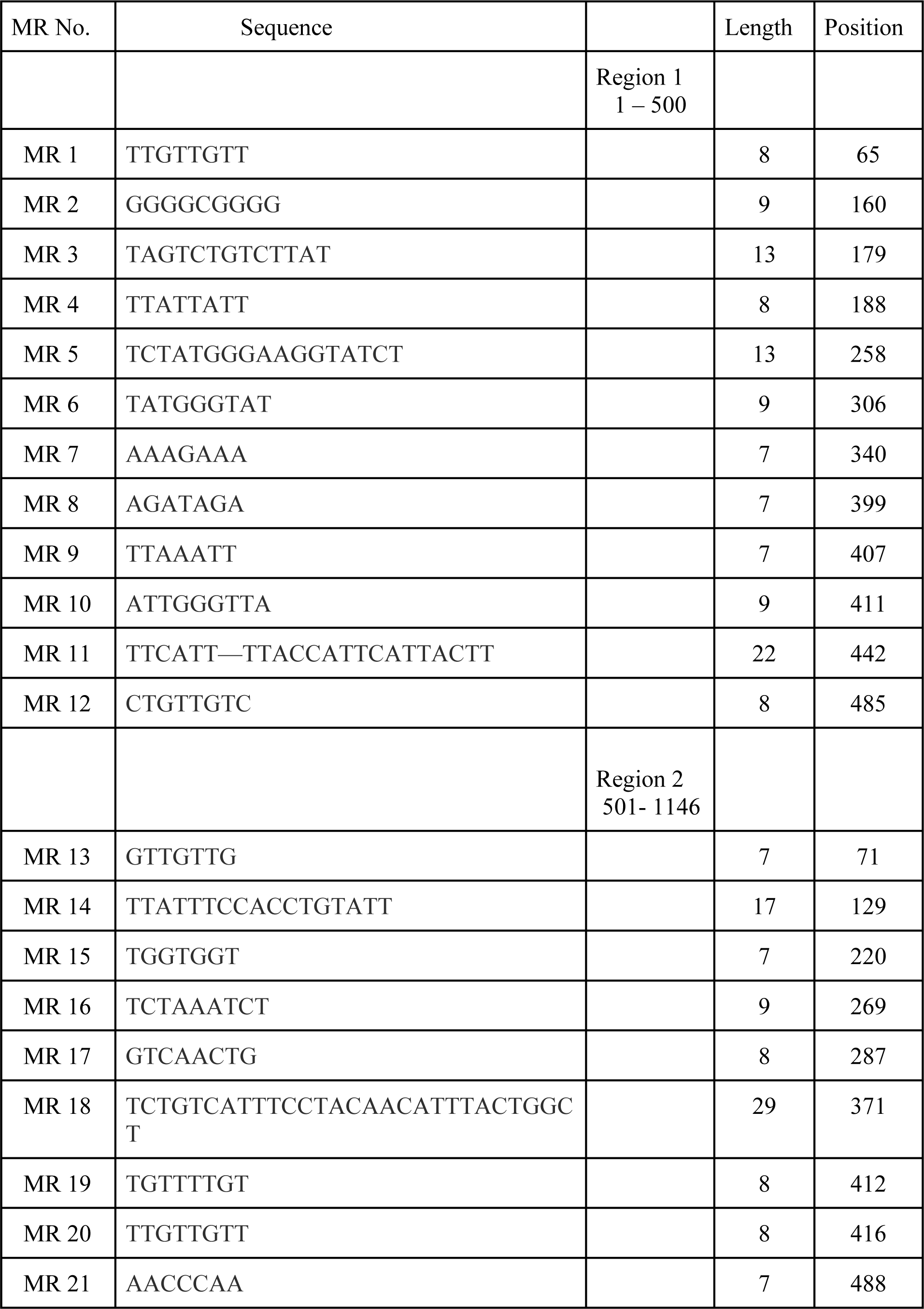
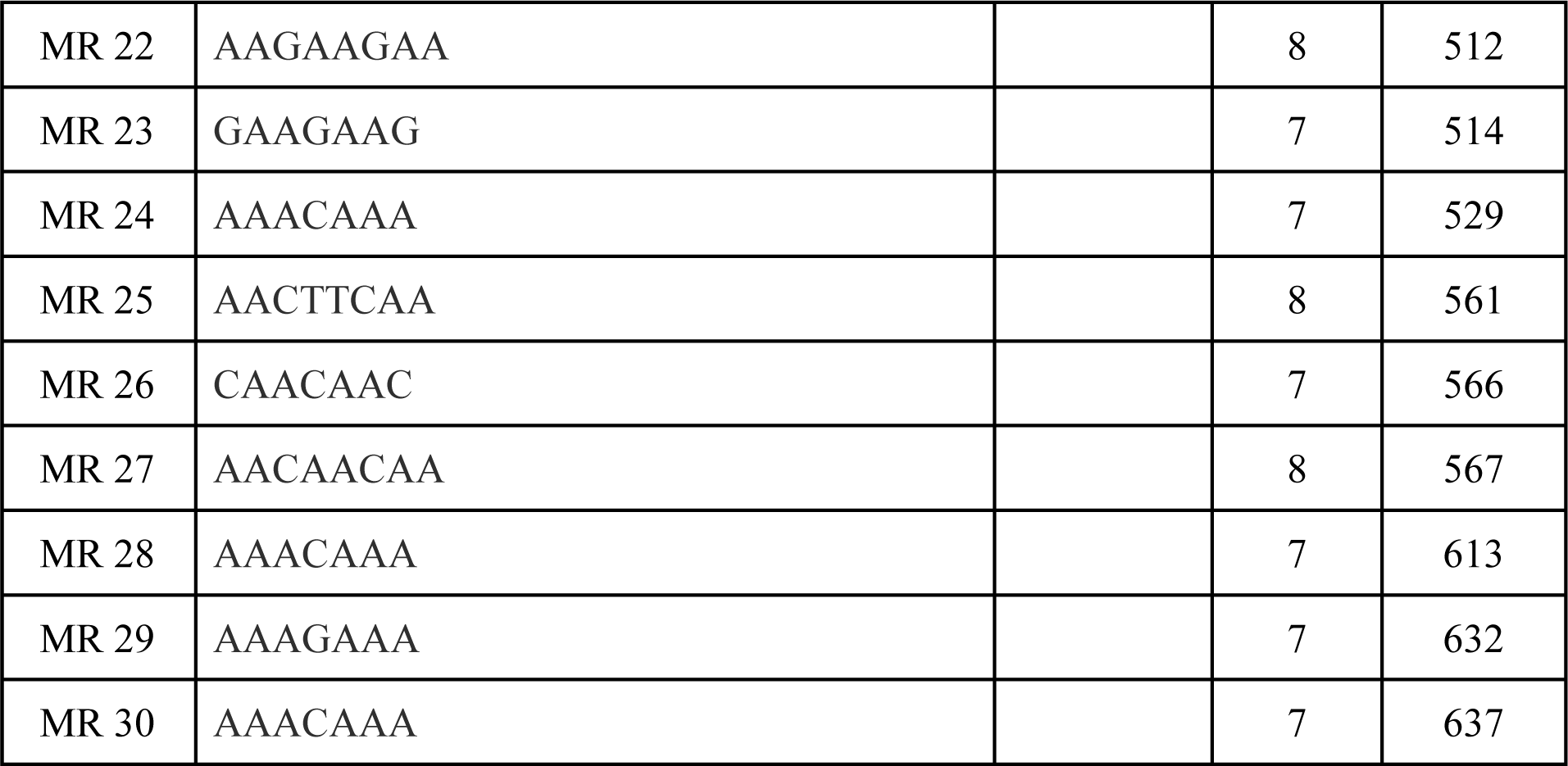
Shows Mirror repeat sequence pattern, their length & position in FTR1 gene.

**Table 5.**
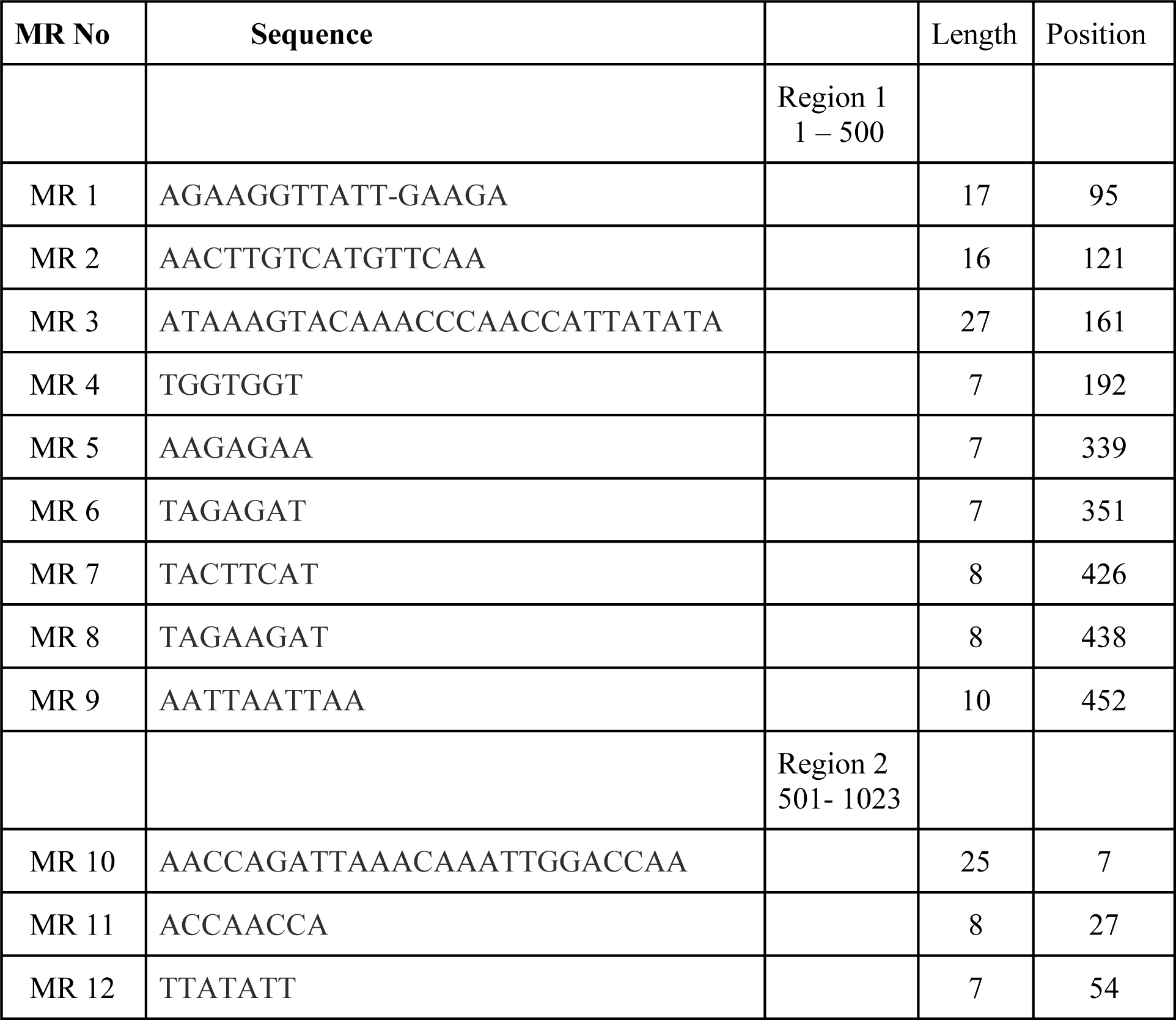
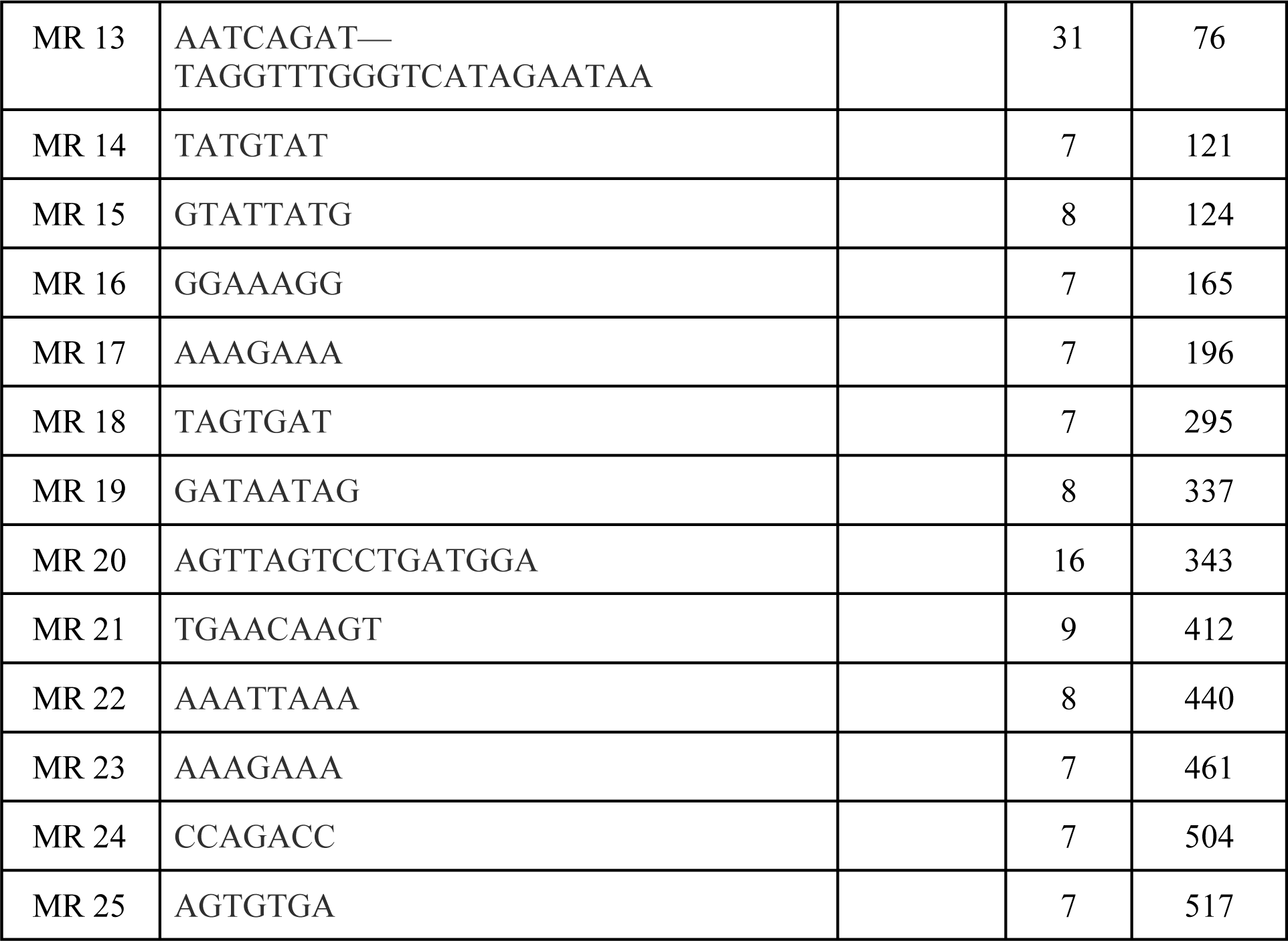
Shows Mirror repeat sequence pattern, their length & position in HEM3 gene.

**Table 6.**
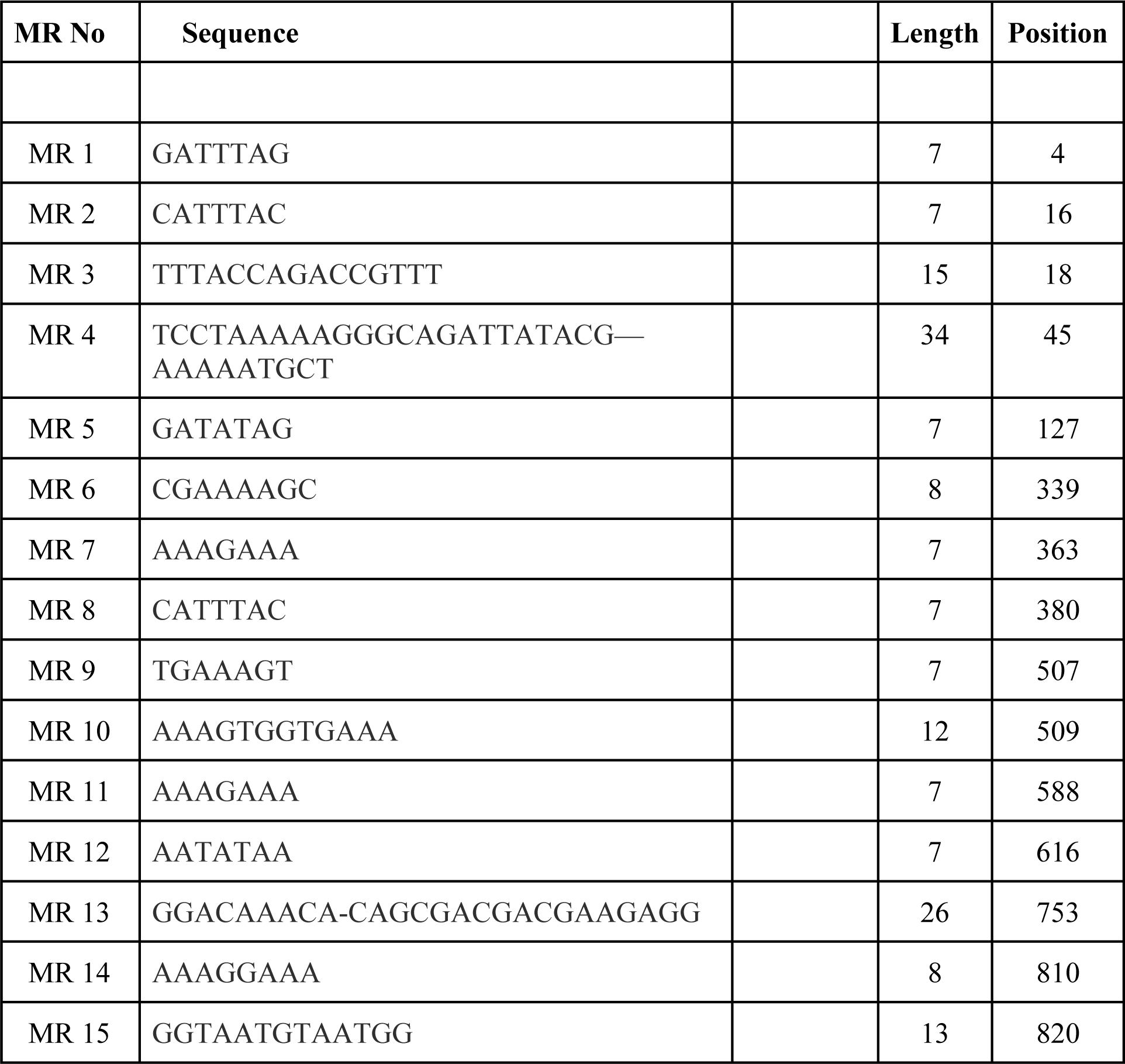
Shows Mirror repeat sequence pattern, their length & position in HIS1 gene.

**Table 7.**
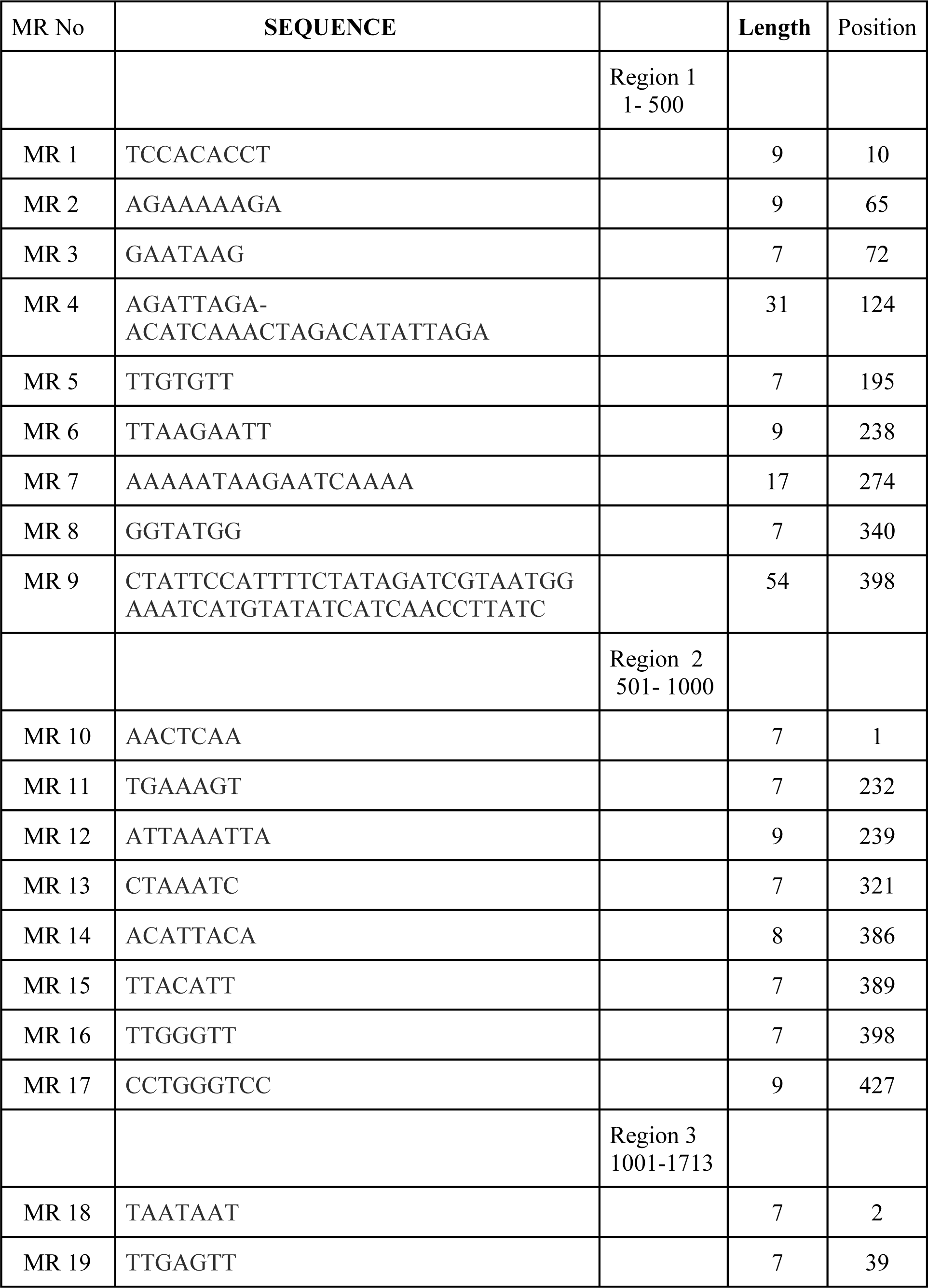
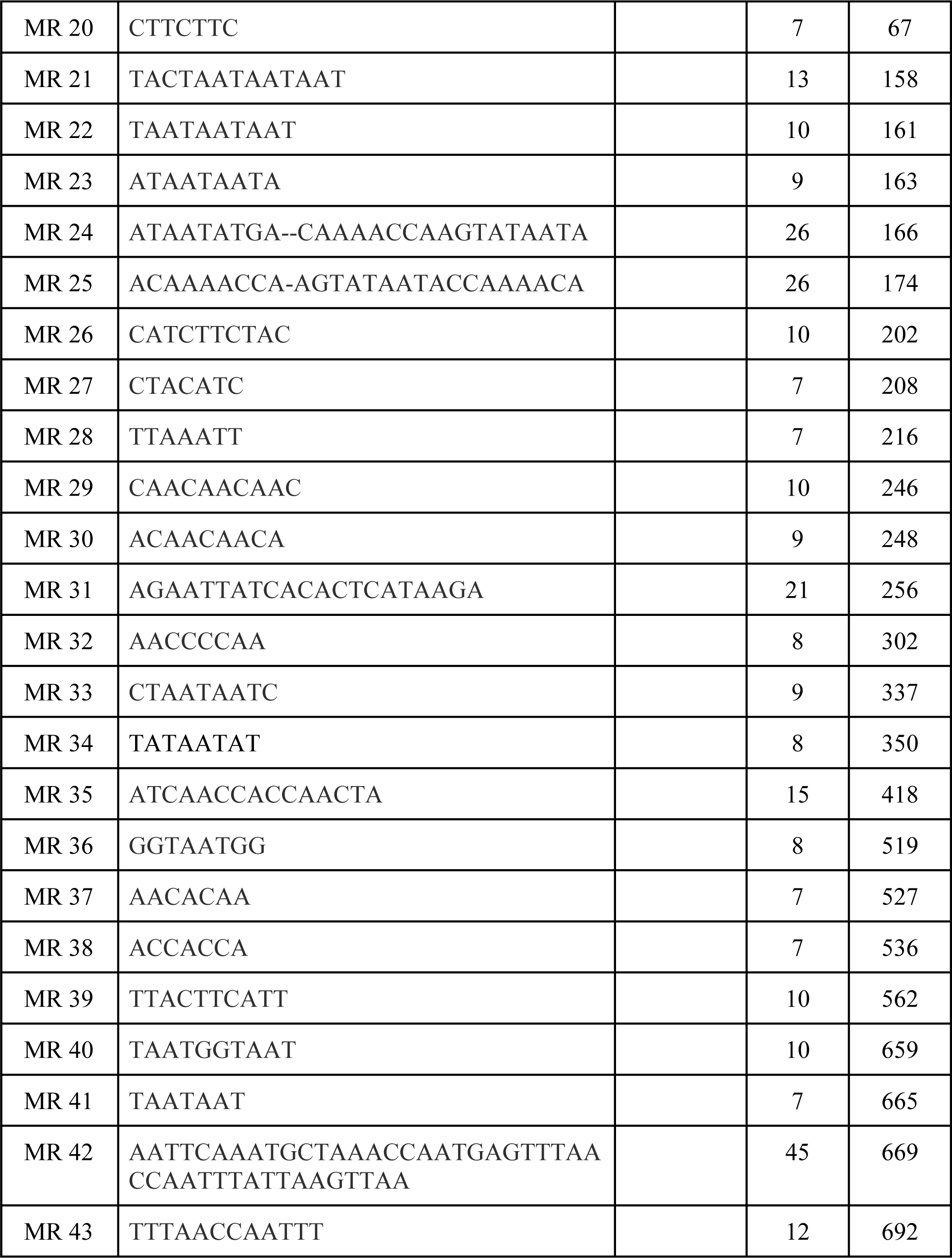
Shows Mirror repeat sequence pattern, their length & position in MIG1 gene.

**Table 8.**
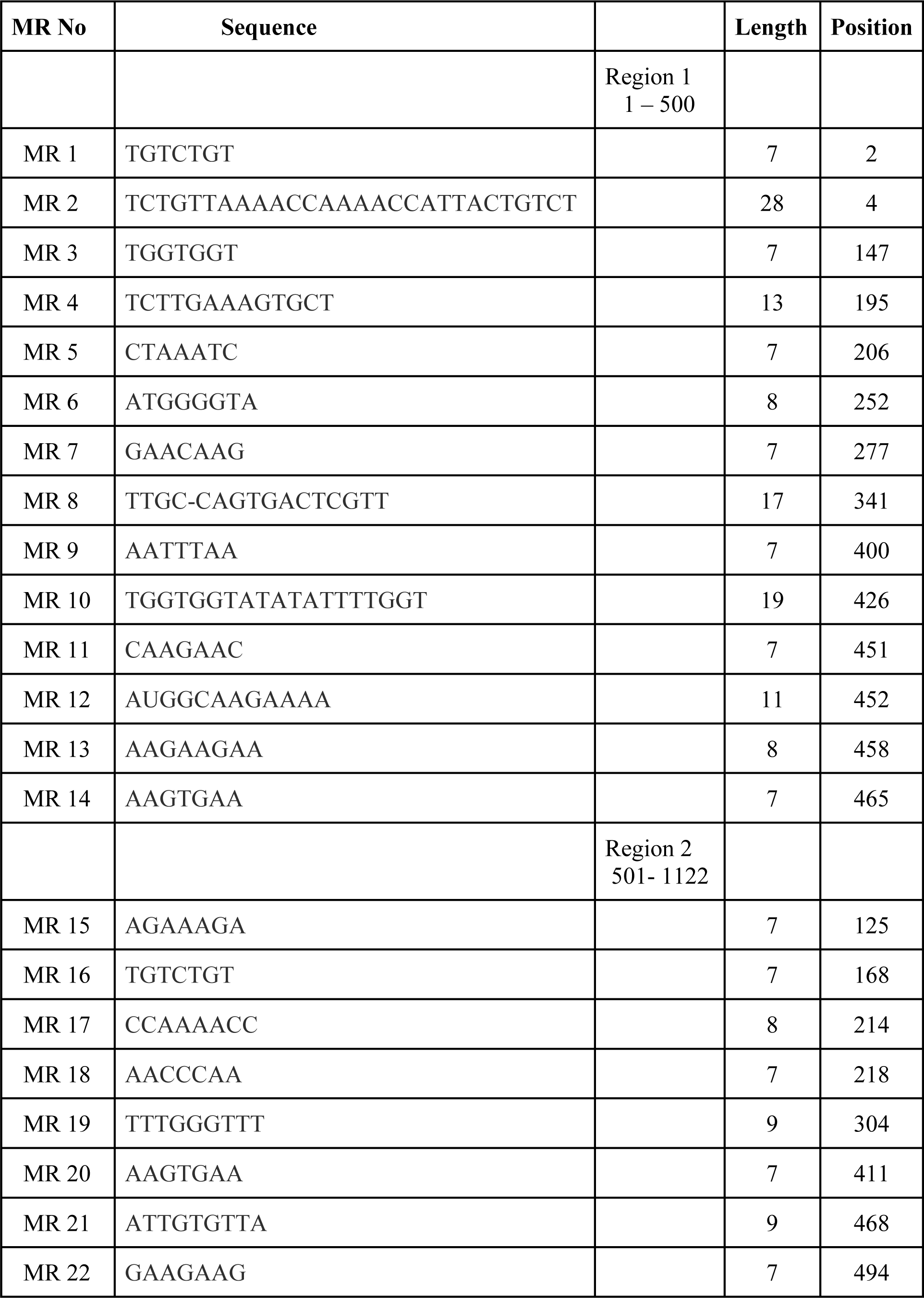
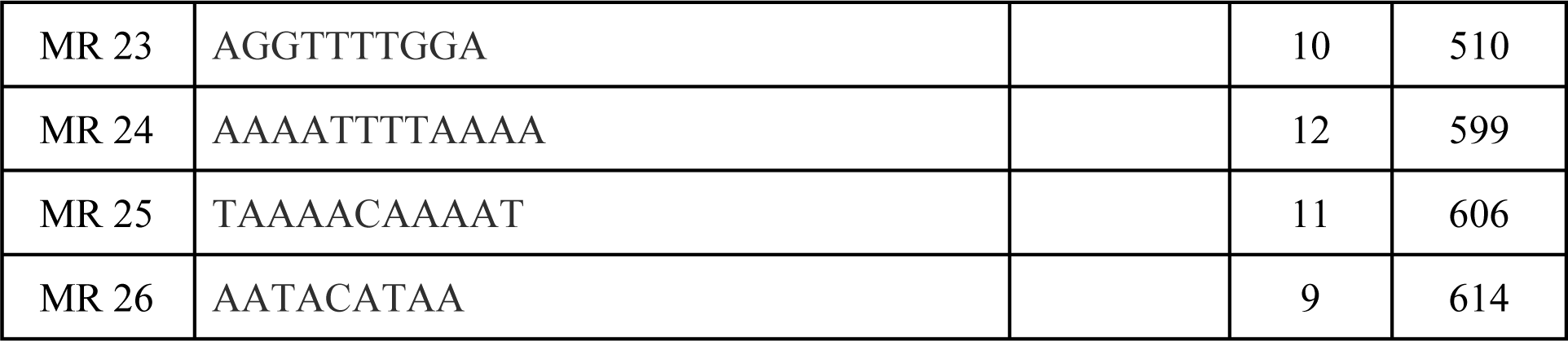
Shows Mirror repeat sequence pattern, their length & position in LEU2 gene.

**Table 9.**
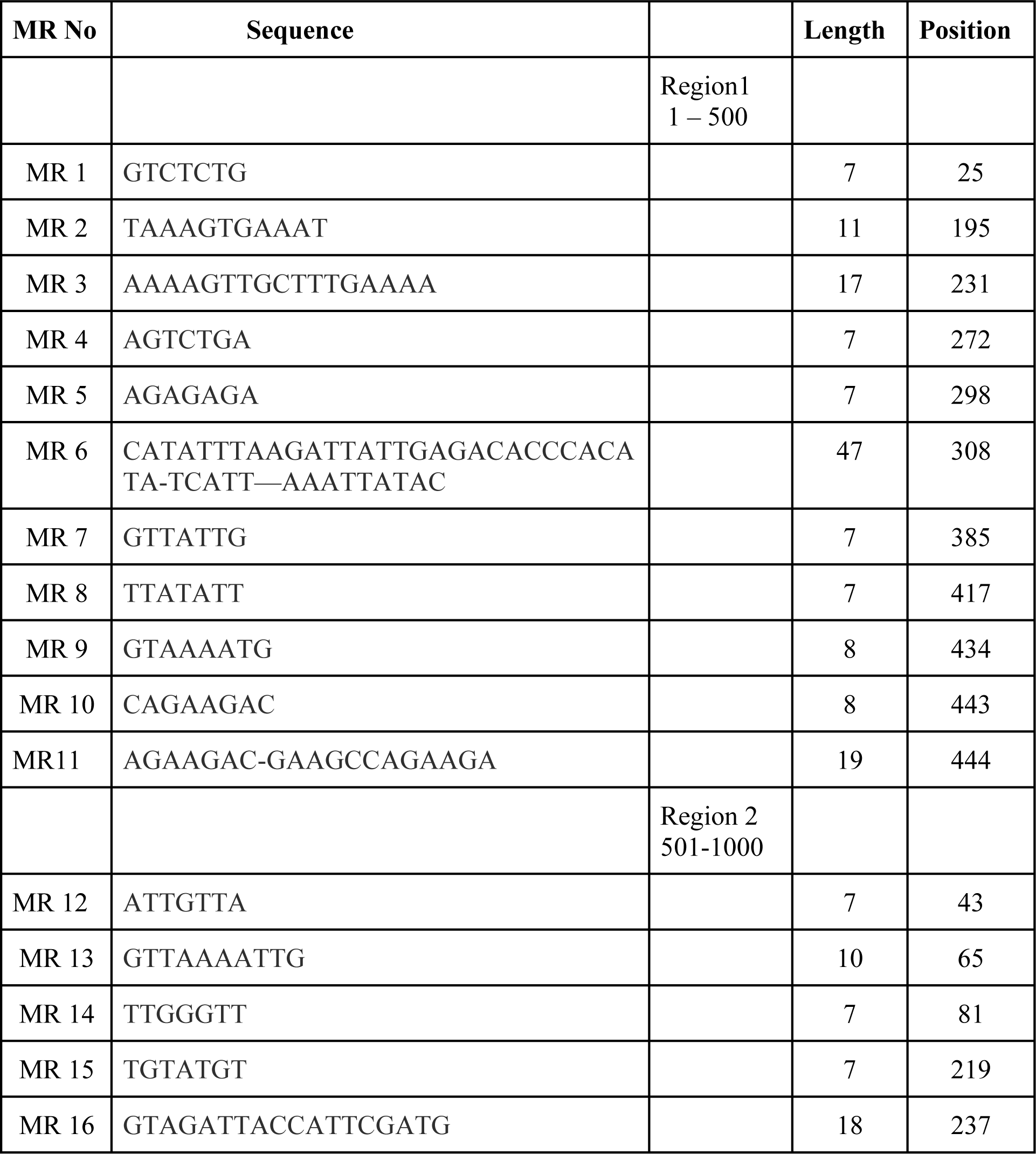
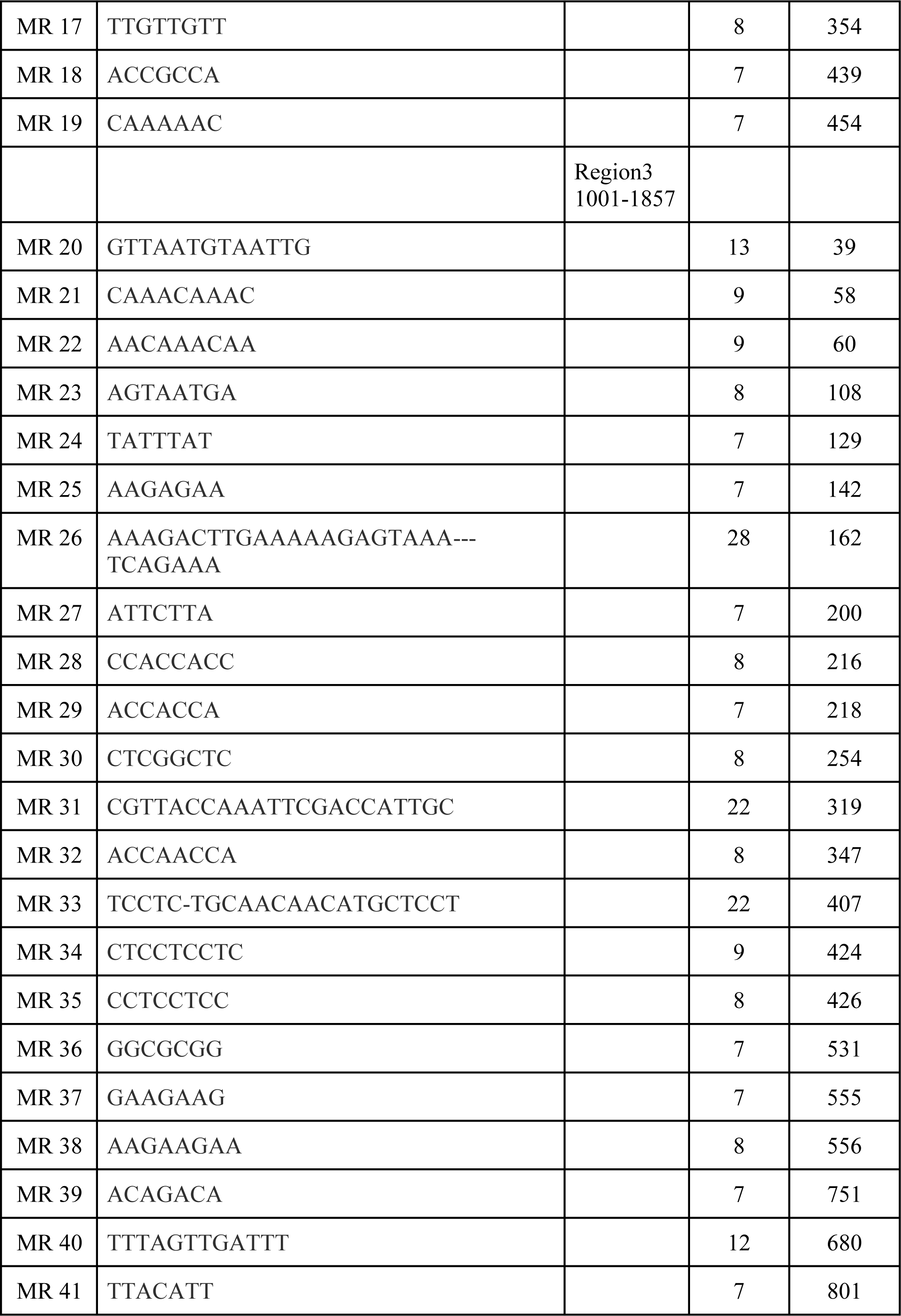
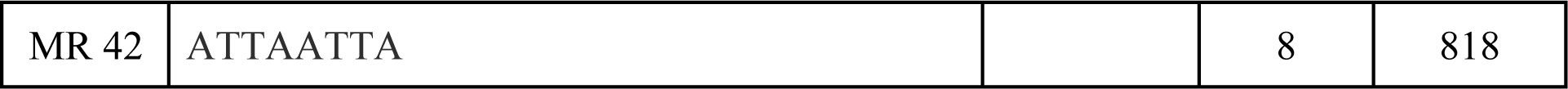
Shows Mirror repeat sequence pattern, their length & position in SNF1 gene.

**Table 10.**
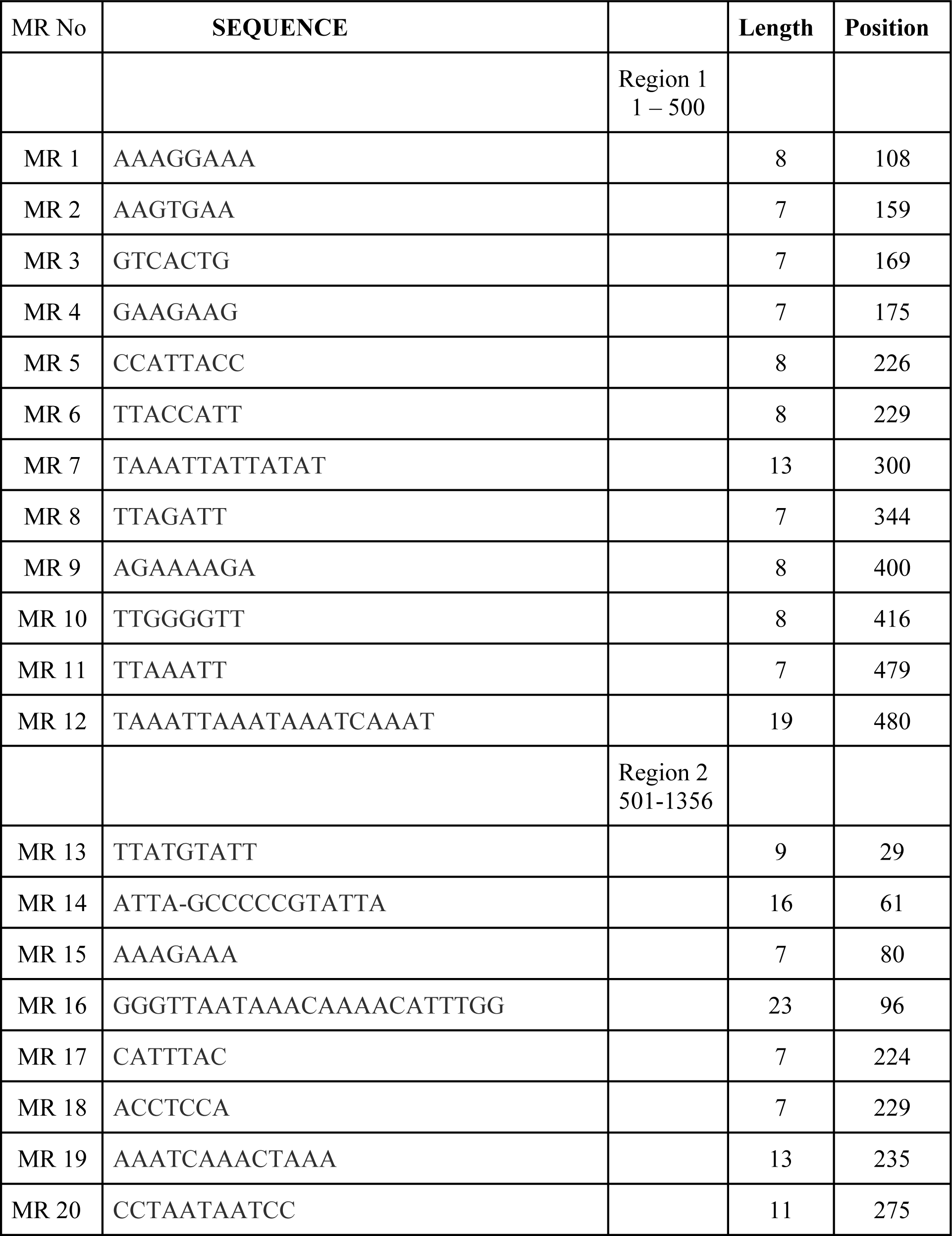
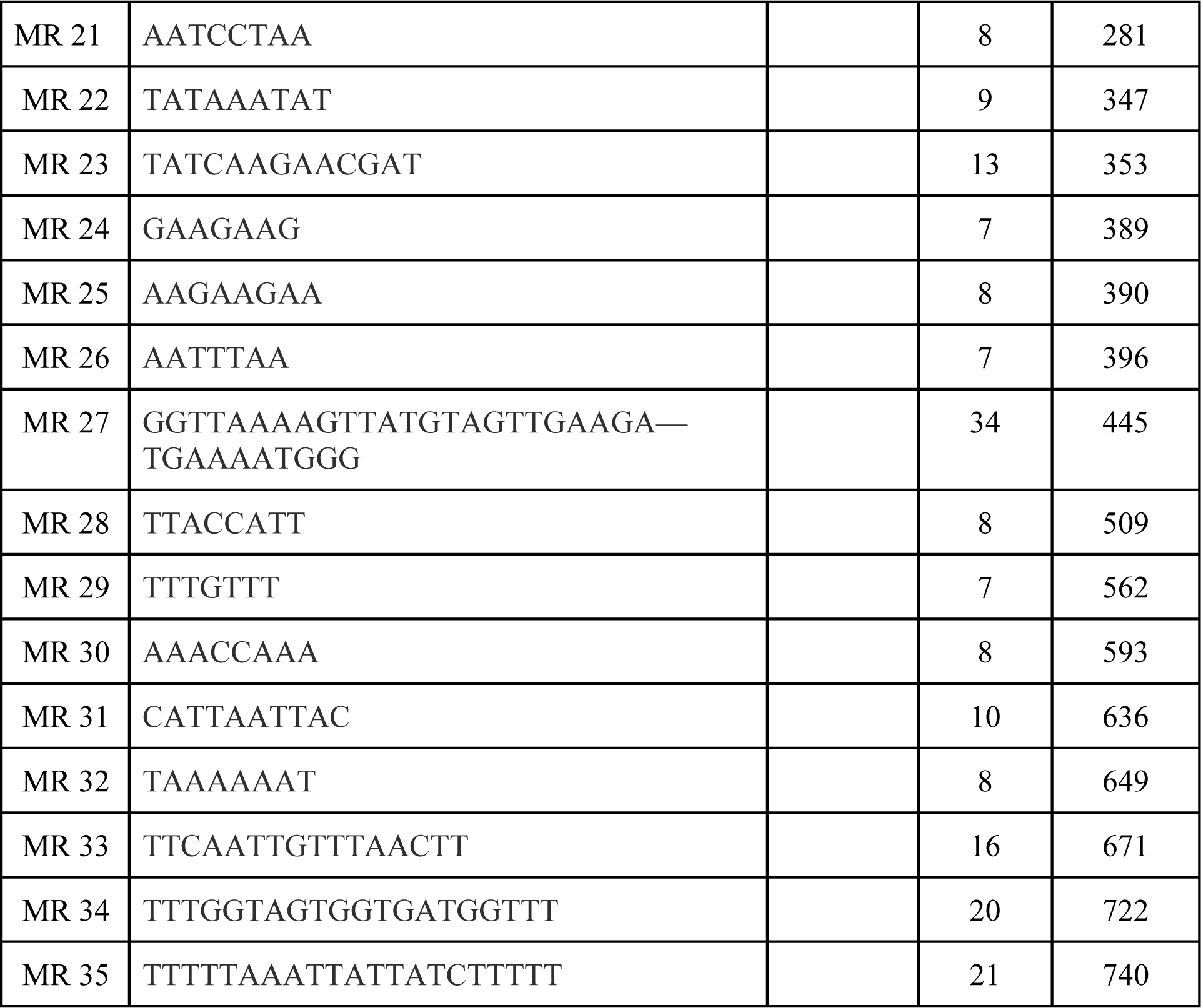
Shows Mirror repeat sequence pattern, their length & position in NMT1 gene.

**Table 11.**
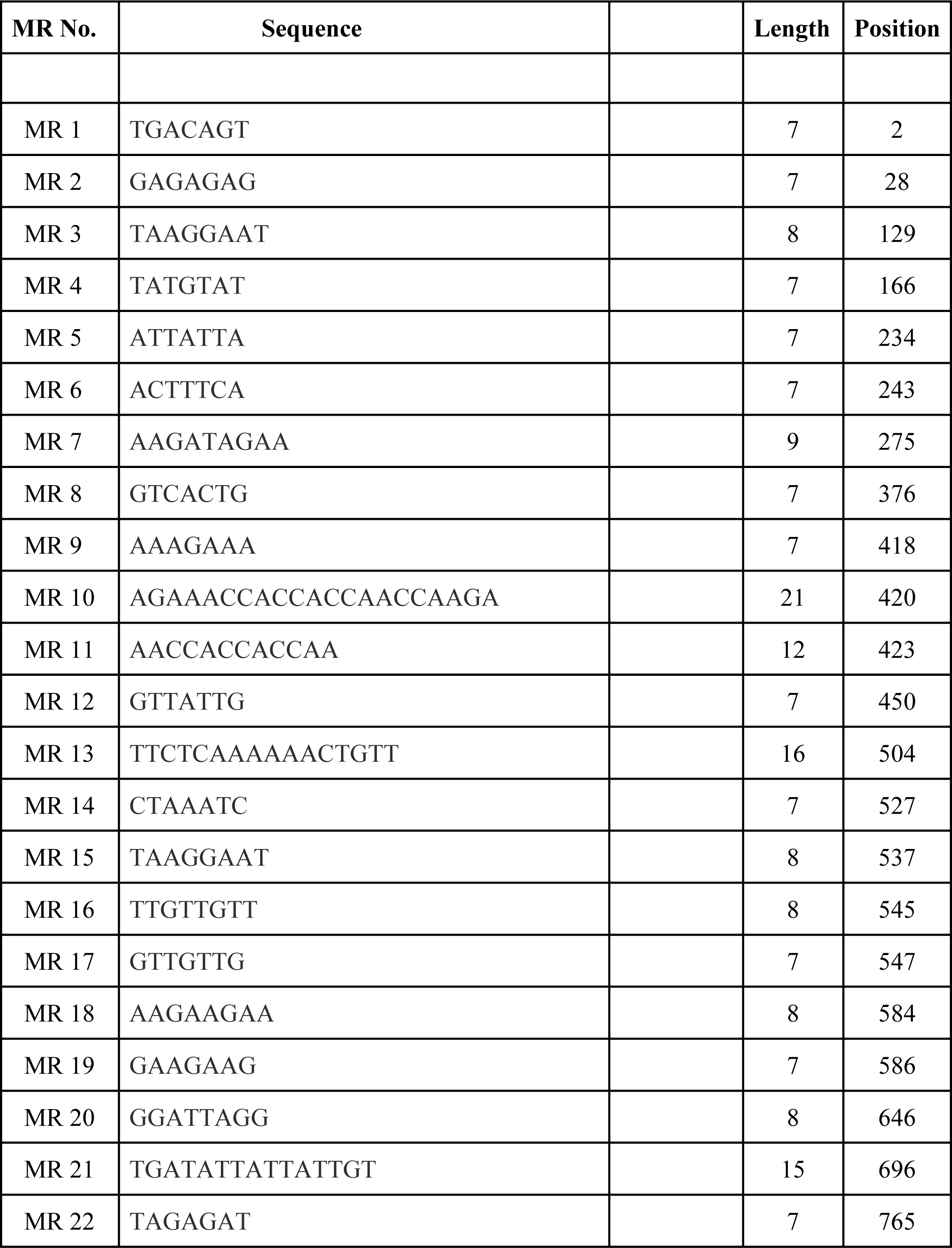
Shows Mirror repeat sequence pattern, their length & position in URA3 gene.

## Discussion

The previous studies on mirror repeats also support the current one. According to the recent study done on human insulin gene were reported a total number of 210 MR sequences in insulin gene (21). Similar like this other studies on animal & plant viruses (19), bacterial genes (16), model plant (17) & some animal domains (24-26) also provide an insight on the frequent occurrence of Mirror repeats in genes/genomes. The output of current study on selected genes of *Candida albicans* also shows that MR sequences frequently present in all the selected genes. These sequences show diversity at the level of their length as well as provide a hint on their specific occurrence in a particular domain. Due to their unique nature of occurrence they will be utilized for many prospective like as a biomarker for testing, in evolution based studies, in context of clinical uses (27) etc. The modern era of computational biology can be used them for phylogeny prospective or to classify the domains due to their conservative nature.

## Conclusion

*In-silico* analysis of selected genes of *Candida albicans* concluded that the frequent occurrence of mirror sequences gives many insights like these sequences will be the integral part of the genes & will be play crucial roles at genetic level in the said fungus. These sequences will be utilized for many purposes like in evolutionary studies, clinical & therapeutic based studies as well as in computational genomics or proteomics based research.

## Author contribution statement

The whole experiments & analysis carried out by Ms. Barkha Sehrawat, Ms. Priya Yadav & Mr. Mustak Sarjeet. Ms. Sakshi Yadav, Ms. Vidhi Yadav, Mr. Parvej Alam, Ms. Nupur Goyal & Ms. Meghali Ahlawat helps in data preparation & its interpretation as well as in manuscript drafting & revision. Dr. Sandeep Yadav (Corresponding author) finalized the final version of this manuscript. All authors approved the final version of the manuscript.

## Funding Acknowledgmen

No any financial support received from any funding agency for the current study.

